# Geometric characterization of the HSV-1 glycoprotein B–amyloid *β* interaction in Alzheimer’s disease using Forman-Ricci curvature

**DOI:** 10.64898/2026.08.26.747308

**Authors:** Léa Bou Dagher, Zihan Han, Shenggao Zhou, Tamas Fulop, Mathieu Desroches, Serafim Rodrigues

## Abstract

Alzheimer’s disease is characterized by the accumulation and aggregation of amyloid-*β*(A*β*), but the molecular mechanisms linking environmental and infectious factors to A*β* conformational changes remain incompletely understood. Herpes simplex virus type 1 (HSV-1) has been proposed as a potential contributor to AD pathology, and interactions between the viral glycoprotein B (gB) and A*β* may influence the conformational behaviour of the peptide. Molecular dynamics (MD) simulations provide atomic-scale information on such interactions, but conventional structural descriptors may not fully capture changes in the organization of residue interaction networks. Here, we introduce a graph-geometric framework based on Forman–Ricci curvature to characterize the evolution of residue interaction networks during MD simulations. Each simulation frame is represented as a residue interaction graph based on C*α*–C*α* contacts, and residue-wise curvature profiles are analysed across time. We apply the framework to A*β*1–42 in isolation and in complex with HSV-1 gB. Conventional MD analyses indicate stable association of the simulated complex, favourable interaction energetics, and conformational changes in A*β*, including a transition from *α*-helical structure toward *β*-turn-rich conformations over the simulated timescale. Forman–Ricci curvature reveals pronounced and spatially localized remodelling of the A*β* residue interaction network in the complex, with the strongest changes concentrated in the C-terminal region. These regions also exhibit reduced temporal curvature fluctuations and progressively distinct geometric behaviour throughout the simulation. Hierarchical clustering further identifies cooperative groups of residues with coordinated curvature dynamics, including a prominent C-terminal domain. Together, these results demonstrate that Forman–Ricci curvature provides a complementary description of biomolecular dynamics by capturing changes in the geometric organization of residue interaction networks that are not directly represented by conventional structural descriptors. The framework provides a general computational approach for studying network-level structural remodelling in protein molecular dynamics and offers a quantitative perspective on the conformational consequences of HSV-1 gB–A*β* association.

## Introduction

Alzheimer’s disease (AD) is the most prevalent neurodegenerative disorder and the leading cause of dementia worldwide. It is characterized by progressive cognitive decline accompanied by the accumulation of extracellular amyloid-*β* (A*β*) plaques and intracellular neurofibrillary tangles composed of hyperphosphorylated tau proteins [5]. Despite decades of intensive research, the molecular mechanisms initiating and driving A*β* aggregation remain incompletely understood, and effective disease-modifying therapies are still lacking [10].

The amyloid cascade hypothesis has long served as the dominant framework for explaining AD pathogenesis by proposing that abnormal production and aggregation of A*β* peptides constitute the primary trigger of neuronal dysfunction [3]. However, increasing experimental and clinical evidence indicates that A*β* aggregation alone cannot fully account for the complexity of the disease. In particular, substantial amyloid deposition has been observed in cognitively healthy individuals [4], whereas cognitive decline often correlates poorly with plaque burden. These observations have motivated the development of complementary hypotheses aimed at explaining the origin and progression of Alzheimer’s disease.

Among these, the infectious hypothesis has gained considerable attention over the past decade [6]. Numerous studies have suggested that microbial pathogens may contribute to AD by promoting chronic neuroinflammation or by directly interacting with A*β*. In particular, Herpes Simplex Virus Type 1 (HSV-1) has emerged as one of the strongest candidates. HSV-1 DNA has been detected in post-mortem brains of Alzheimer’s patients, and epidemiological studies have associated recurrent HSV-1 infection with an increased risk of dementia. Furthermore, accumulating experimental evidence suggests that A*β* itself may function as an antimicrobial peptide capable of binding viral particles and promoting their aggregation. Within this framework, viral-induced conformational changes of A*β* may constitute an important step linking infection to amyloid aggregation.

Molecular dynamics (MD) simulations have become an essential tool for investigating protein conformational dynamics at atomic resolution. Classical MD analyses typically characterize structural evolution through descriptors such as the root-mean-square deviation (RMSD), root-mean-square fluctuation (RMSF), hydrogen-bond occupancy, residue contact maps, solvent-accessible surface area, and secondary-structure evolution. These quantities provide valuable information regarding structural stability and local flexibility. Nevertheless, they primarily describe individual geometric or energetic properties and often overlook the global organization of residue interaction networks that emerge during protein motion.

Graph theory offers a natural framework for representing biomolecular structures as interaction networks. In such representations, residues are modelled as graph vertices while spatial contacts define the edges. This perspective enables the application of discrete geometric tools that characterize not only individual residues but also the organization of their interactions. Among these tools, discrete Ricci curvature has recently attracted increasing interest in network science because it quantifies the local geometry of graphs while simultaneously capturing global structural organization. In biological systems, Ricci curvature has been successfully applied to the analysis of protein structures, interaction networks, and molecular docking, where it provides information that is complementary to conventional structural descriptors.

In this work, we introduce a graph-geometric framework for analysing molecular dynamics trajectories based on Forman–Ricci curvature [2]. Each molecular dynamics frame is converted into a residue interaction graph constructed from C*α* atoms, and the Forman–Ricci curvature is computed for every residue throughout the trajectory. This procedure transforms a molecular dynamics simulation into a temporal sequence of discrete geometric signatures, allowing structural evolution to be investigated from the perspective of graph geometry rather than solely through atomic coordinates.

We apply this framework to molecular dynamics trajectories of the A*β* monomer and the HSV-1 glycoprotein B (gB)–A*β* complex. By comparing the curvature dynamics of the two systems, we investigate whether viral binding induces measurable changes in the geometric organization of residue interactions. Several complementary analyses are performed, including comparisons of average curvature profiles, temporal fluctuations, residue-specific curvature trajectories, and spatio-temporal curvature maps.

Our results demonstrate that Forman–Ricci curvature captures localized and persistent geometric remodelling associated with HSV-1 binding. In particular, the largest geometric changes consistently occur within the aggregation-prone C-terminal region of A*β*, while the overall residue interaction network becomes dynamically more stable throughout the simulation. These findings illustrate how discrete differential geometry provides a complementary perspective on protein conformational dynamics and establish graph Ricci curvature as a promising descriptor for the analysis of molecular dynamics trajectories.

## Materials and methods

### Structure preparation

#### Structure for HSV-1 gB–A*β***_1_*_−_*_42_** complex

First of all, molecular docking was performed using Rosetta (version 2024.09) to derive the HSV-1 gB– A*β*_1*−*42_ complex. The structures of full length A*β*_1*−*42_ (PDB ID: 1IYT) and the HSV-1 gB (PDB ID:8KFA) were retrieved from the protein data bank (PDB). Rosetta is a comprehensive software suite for modelling macromolecular structures based on Monte Carlo simulated annealing, which is widely used for remodelling of proteins and nucleic acids. In our study, we performed protein-protein docking using local docking protocol, followed by structure relaxation with relax module. The generated decoys were clustered using cluster module. The final binding structure with the best score was selected based on the most populated cluster.

#### Structure for individual simulation of HSV-1 gB–A*β***_1_*_−_*_42_** complex

The structure of A*β*_1*−*42_ for individual simulation is identical to the one used in the complex. The structure for HSV-1 gB has been replaced with AlphaFold structure because of a long string of missing residues in 8KFA and the other structures in PDB, which caused GROMACS to pull the residues at both ends together, leading to fragmentation, as shown in Figure 1) and killing the simulation process. The comparison between 8KFA and AlphaFold structure is shown in Figure 2

**Figure 1.**
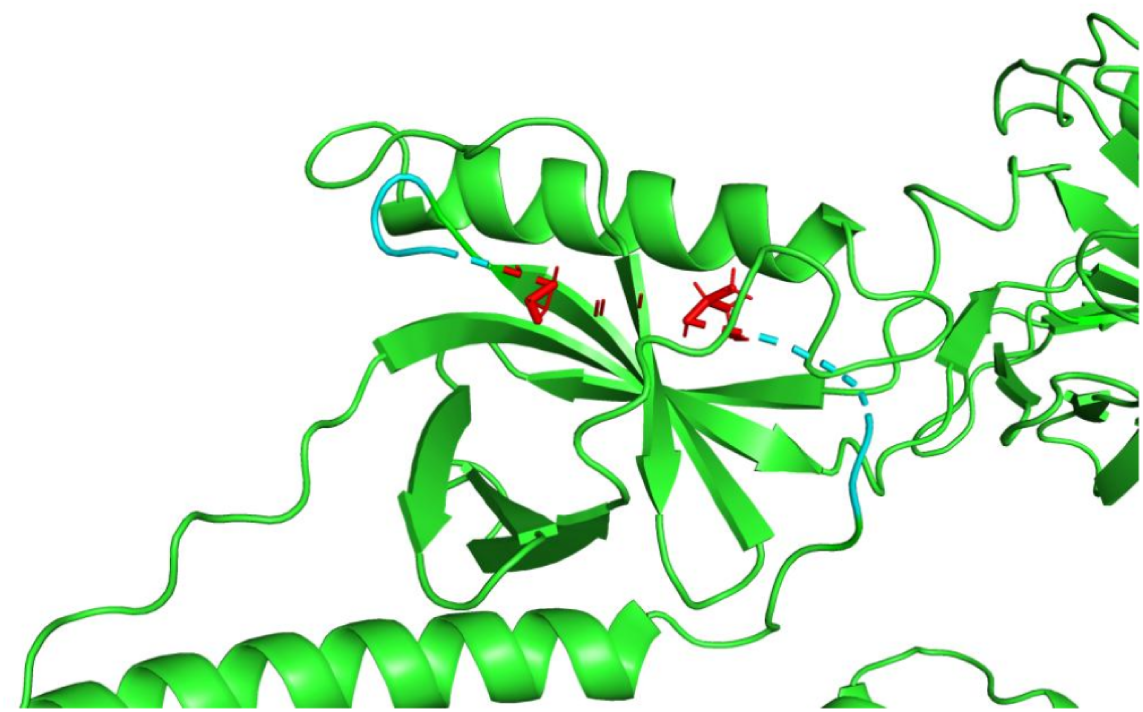
The fragmentation occurred in the simulation of HSV-1 gB (PDB ID:8KFA). The fractured residues are coloured in red which are two ends of the string of missing residues, and the three closest residues of each end are coloured in cyan.

**Figure 2.**
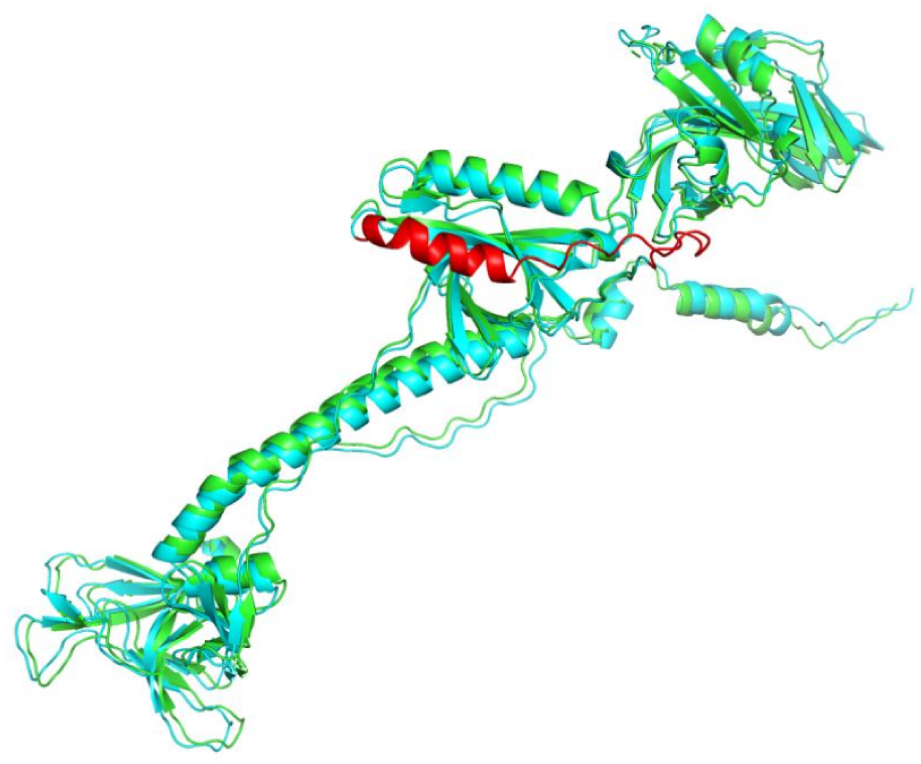
The comparison between 8KFA (coloured in green) and AlphaFold structure of HSV-1 gB (coloured in cyan and mended parts coloured in red).

### Molecular dynamics (MD) simulations

The predicted conformation of the HSV-1 gB–A*β*_1*−*42_ complex and each protein were further refined by performing three individual MD simulations using GROMACS 2021.7 software with the Amber99sb force field. For each simulation, the initial structure was solvated in a rectangular box of the simple point charge (SPC) water model, and the system in the solvent box was neutralized with appropriate amount of Na+ ions. Before the MD simulation, the solvated system was fully energy-minimized with the steepest descent and conjugate gradient methods. Then, the system was equilibrated by the processes 100 ps at 298.15 K with NVT ensemble and 100 ps at 1 bar with NPT ensemble prior to a sufficiently long MD simulation. The 500 ns explicit MD simulation was performed with a periodic boundary condition in the NPT ensemble using a Parrinello-Rahman ensemble with a pressure(P) of 1 bar, and a V-rescale thermostat at a temperature of 298.15 K.

### Evaluation of binding free energy

The binding free energy between HSV-1 gB–A and *β*_1_*_−_*_42_ was evaluated by using molecular mechanics generalized Born surface area method (MM/GBSA) through gmx MMPBSA of GROMACS.

### Residue interaction graph construction

Each molecular dynamics frame is represented by an undirected graph

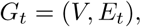

where *t* denotes the simulation time.

Each vertex *v_i_ ∈ V*, corresponds to the C*α* atom of one amino-acid residue. Since the peptide contains 42 residues, every graph consists of 42 vertices, while only the edge set evolves during the simulation. Two residues are considered to interact whenever the Euclidean distance between their C*α* atoms is smaller than a prescribed cut-off distance *d_c_*= 1 nm. Consequently, an edge

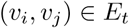

is introduced whenever

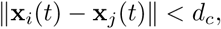

where **x***_i_*(*t*) denotes the Cartesian coordinates of residue *i* at time *t*.

This construction transforms every molecular configuration into a residue interaction network that captures the instantaneous spatial organization of the protein.

### Forman–Ricci curvature

To quantify the local geometry of each residue interaction graph, we employ the Forman–Ricci curvature, a discrete analogue of Ricci curvature originally introduced in discrete differential geometry.

For an unweighted graph, the Forman curvature associated with an edge *e* = (*u, v*) is given by

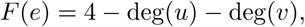

where deg(*u*) and deg(*v*) denote the degrees of the two incident vertices.

The GraphRicciCurvature Python package subsequently assigns a curvature value to every vertex by combining the curvature values of all incident edges. This vertex curvature provides a quantitative measure of the local geometric environment surrounding each residue within the interaction network.

Unlike classical structural descriptors that focus on individual distances or energies, Forman–Ricci curvature simultaneously reflects local connectivity, network organization and the structural role of each residue inside the evolving interaction graph.

### Graph-Based analysis pipeline

For every frame of the molecular dynamics trajectory, the C*α* coordinates are first extracted to construct the corresponding residue interaction graph. Forman–Ricci curvature is then computed for every residue, yielding a curvature vector describing the geometric state of the protein at that time point.

Repeating this procedure for every frame produces a curvature matrix whose rows correspond to simulation time and whose columns correspond to amino-acid residues. This matrix forms the basis of all subsequent analyses.

Four complementary analyses are performed. First, average curvature profiles are compared to identify residues undergoing persistent geometric remodelling upon HSV-1 binding. Second, temporal fluctuations of residue curvature are analysed to evaluate changes in structural stability. Third, residue-specific curvature trajectories are investigated to characterize the temporal evolution of the most strongly affected regions. Finally, spatio-temporal curvature maps and clustering analyses are employed to identify groups of residues exhibiting similar geometric behaviour throughout the simulations.

## Results

Before investigating the geometric organization of the residue interaction network, we first present the conventional molecular dynamics analyses of the HSV-1 gB–A*β* complex. These analyses establish the structural stability, intermolecular interactions, and conformational behaviour of the complex throughout the simulations, providing the structural context for the graph-based analyses that follow. We then examine the same trajectories using Forman–Ricci curvature to determine whether the residue interaction network reveals complementary geometric signatures associated with HSV-1 binding.

### Molecular docking of the HSV-1 gB–A*β*_1_*_−_*_42_ complex

The initial binding mode between A*β*_1*−*42_ and HSV-1 gB was theoretically predicted by molecular docking method. Firstly, we conducted a brief literature review to obtain knowledge of the reported binding region of HSV-1 gB. Fortunately, Sun et al has revealed the structure of HSV-1 gB bound to D48 in DII domain [7] (see Figure 3). On this basis, we implemented a DII domain-specific local docking protocol. According to the docking score and cluster analysis, we got the most potential binding structure which the total score is -1232.879 REU and interface score is -17.484 REU. Then, the folding of the structure was monitored through 500 ns MD simulation.

**Figure 3.**
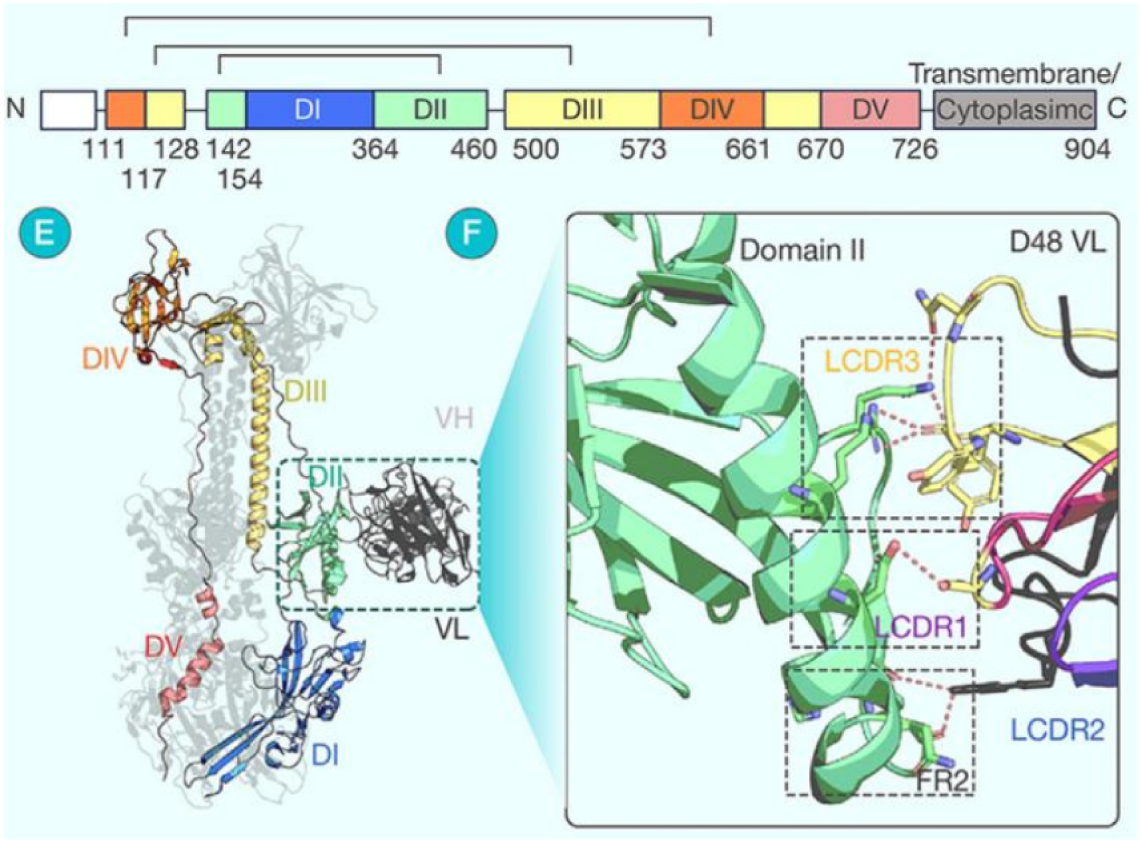
The structure domain of HSV-1-gB [7].

**Figure 4.**
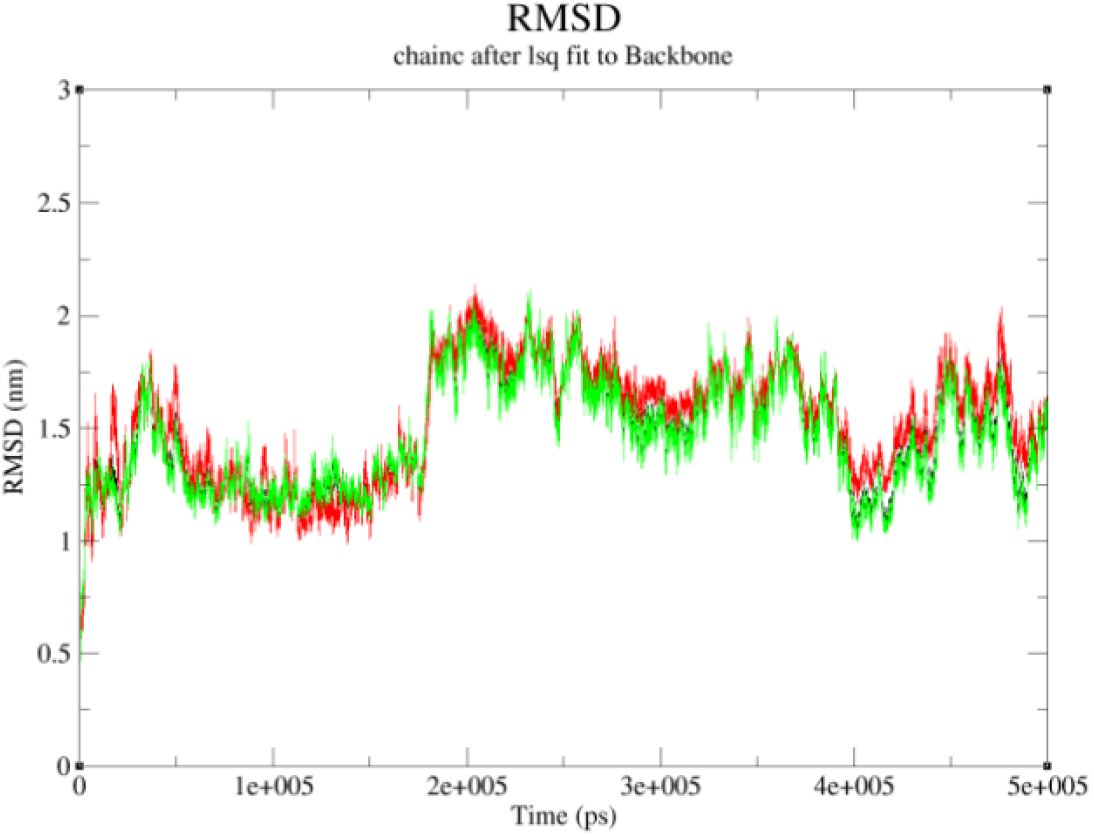
The RMSD of full structure of the HSV-1 gB – A*β*_1*−*42_ complex. The RMSD of A*β*_1*−*42_ is coloured in red, HSV-1 gB in green and complex in black.

### RMSD analyses

The root mean square deviation (RMSD) is considered as the important parameter to assess the structural stability during the MD simulation. At first, we use the full structure of A*β*_1*−*42_ and HSV-1 gB which we consistently observed conformational fluctuations even though the much longer simulation than the MD simulation performed in [8] (see Figure **??**). Then, trajectory snapshots were extracted at 10 ns intervals from the last 100 ns of the simulation (see Figure 5). From the snapshots, we found that the binding domain between A*β*_1*−*42_ and HSV-1 has actually been stable and the part of HSV-1 gB which we think is away from the binding domain is fluctuating. As a result, we extracted the structure which nearby the binding domain for further analysis and the analyses of the MD trajectory showed that the RMSD of HSV-1 gB – A*β*_1*−*42_ complex started to be stable after 250 ns, with RMSD values at 1.06 *±* 0.05 nm, 1.02 *±* 0.05 nm and 1.60 *±* 0.05 nm, respectively (see Figure 6).

**Figure 5.**
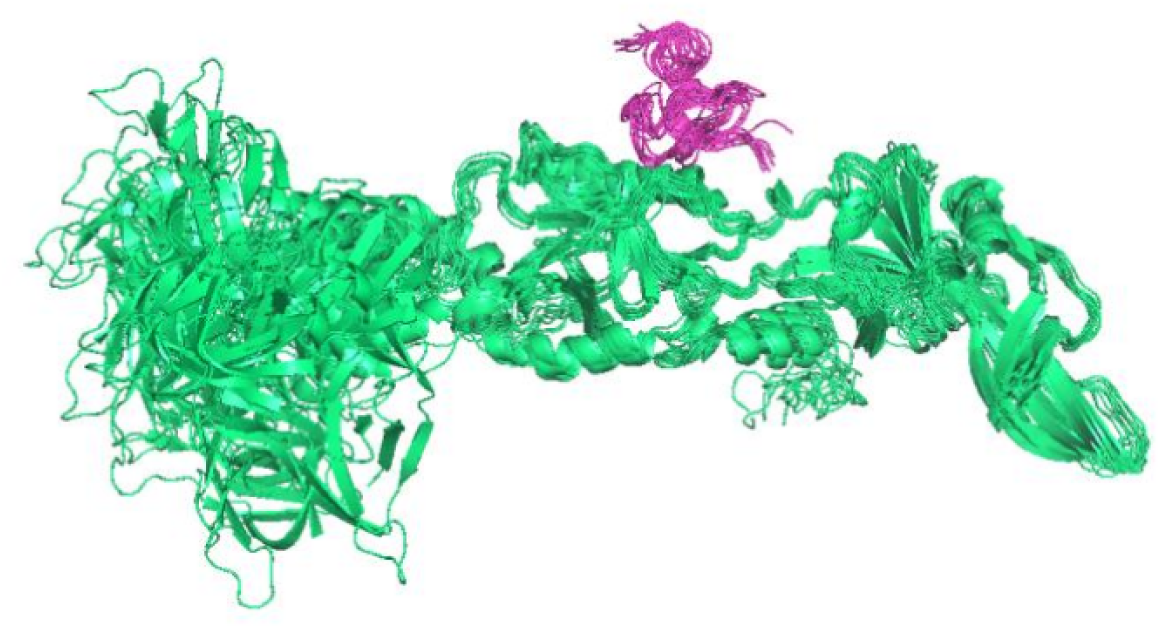
The snapshots from last 100 ns of the simulation. A*β*_1*−*42_ is coloured in light magenta and HSV-1 gB is coloured in lime green.

**Figure 6.**
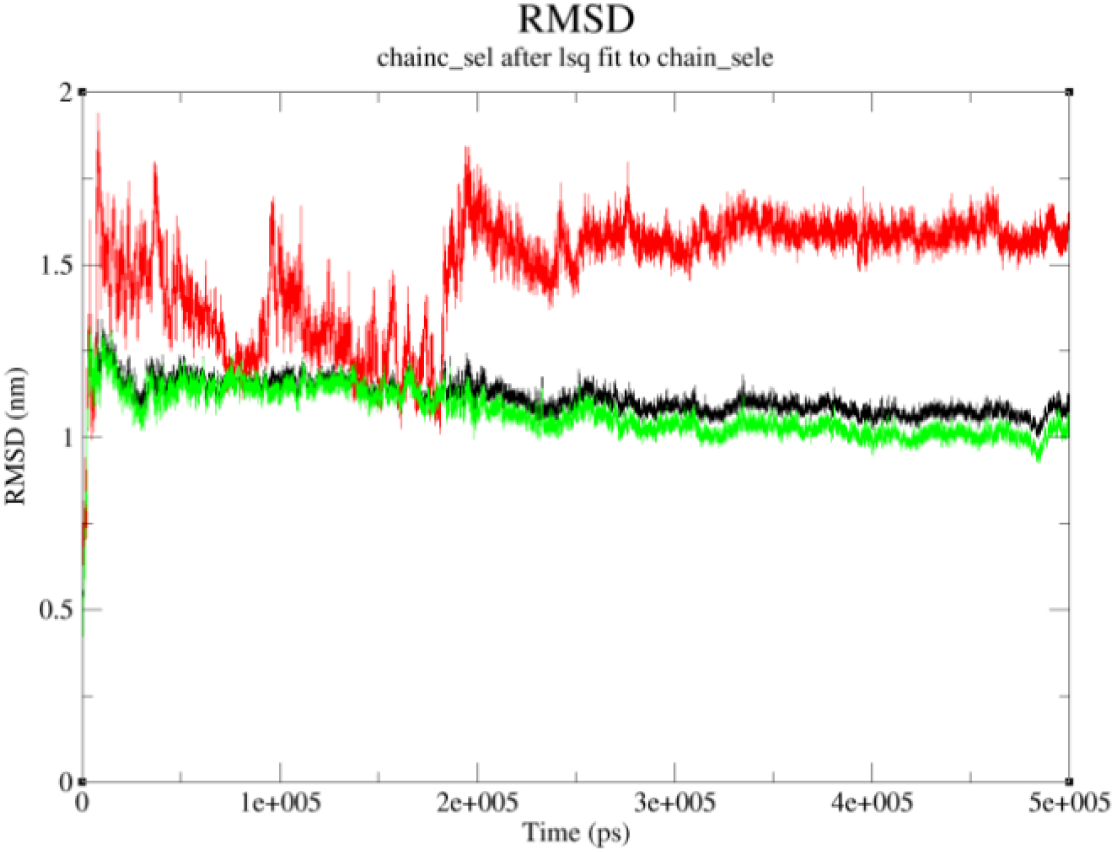
The RMSD of binding domain of the HSV-1 gB - A*β*1-42 complex. The RMSD of A*β*1-42 is coloured in red, HSV-1 gB in green and complex in black.

### Conformations and binding mode analysis of the HSV-1 gB–A***β*_1_*_−_*_42_** complex

From MD trajectories, we found that the conformational transition is pronounced, especially in AA*β*_1*−*42_. As shown in Figure 7(A, B), the conformational transition sequentially from *α*-helix to a disordered structure, and then to *β*-turn in our simulation, which is consistent with the result in [9] shown in Figure 7(C). This result is different from the conformations observed in [8], which ran the 50ns MD simulation of HSV-1 gD-AA*β*_1*−*42_ complex in water solution with GROMOS 96 53a7 force field. Hence, we conducted a brief literature review[4–6] and found that the conformation of AA*β*_1*−*42_ has a tendency to transform from *α*-helix to *β*-sheet from residue 17 to residue 42 in solvent. The process of the conformational conversion also represents the initial phase of fibrillisation which is one of the important pathogenic mechanisms in Alzheimer’s disease. (More information can be found in 3. Structure analysis part)

**Figure 7.**
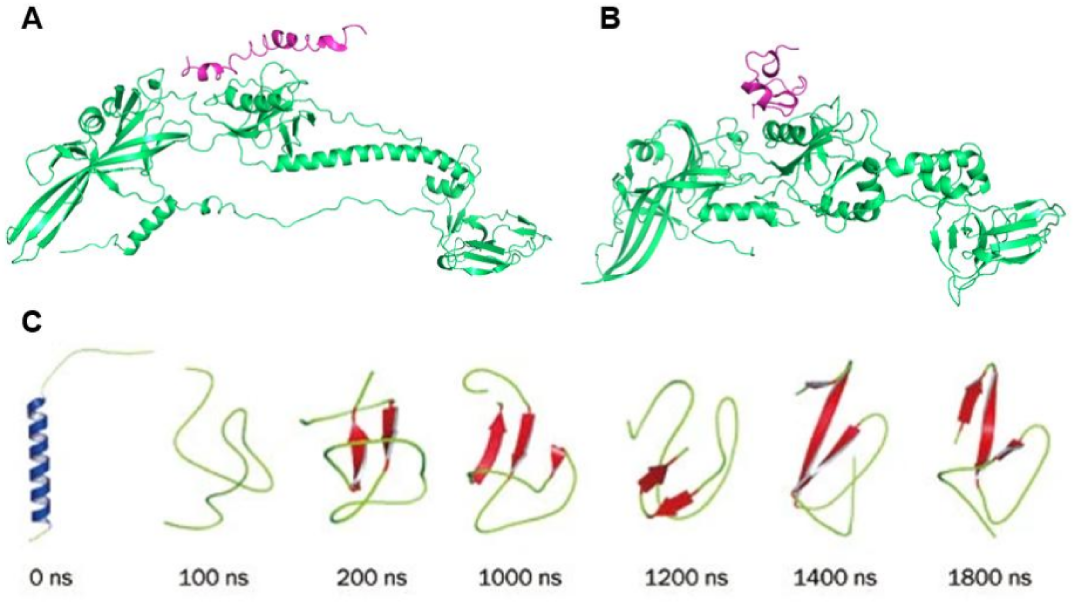
The initial conformation (A) and the stable conformation (B) of the MD trajectory. The conformational conversion of A*β*_1*−*42_ in solution (C) [9].

Then, we visualized the interactions between HSV-1 gB and A*β*_1*−*42_ using PyMol v2.4.1. As shown in Figure 8, there are six hydrogen bonds formed between HSV-1 gB and A*β*_1*−*42_ in range of 0.16-0.28 nm. The formations of these hydrogen bonds effectively maintain the stability of the HSV-1 gB - A*β*_1*−*42_ complex.

**Figure 8.**
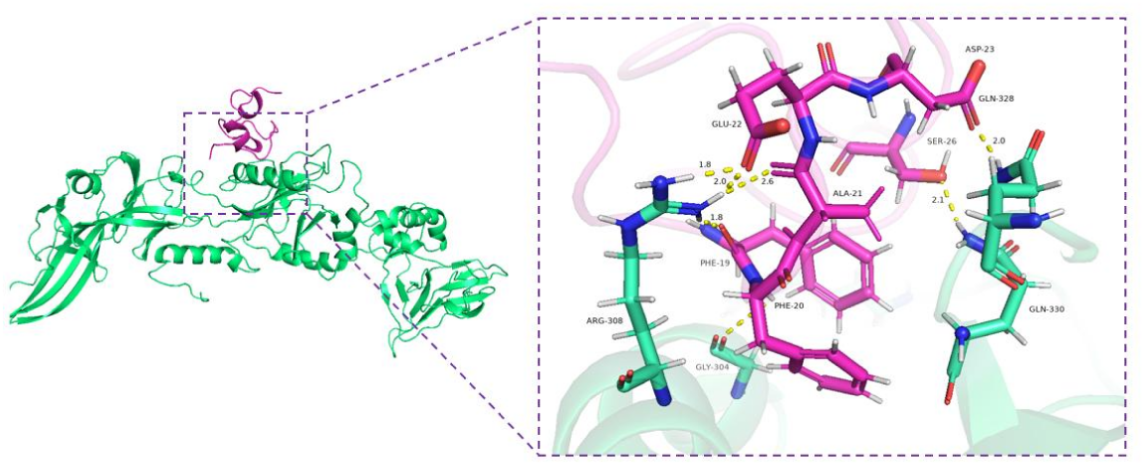
The binding mode of the HSV-1 gB - A*β*_1*−*42_ complex. A*β*_1*−*42_ is coloured in light magenta and HSV-1 gB is coloured in lime green. The hydrogen bonds formed between chains are denoted in yellow dash.

### Quantitative analyses of the binding mode for the HSV-1 gB –A***β*_1_*_−_*_42_** complex

In order to get a quantitative insight into the binding mode of HSV-1 gB–A*β*_1*−*42_ complex, the binding free energy of the complexes was decomposed using the gmx MMPBSA method. The binding free energy was calculated to be -34.49 kcal/mol, and van der Waals (Δ*E_vdw_*) interaction and electrostatic (Δ*E_ele_*) interaction are the favourable components of the binding affinity, -50.70 kcal/mol and -37.64 kcal/mol respectively. According to the binding energy decomposition, the main contributions to the binding stability between HSV-1 gB and A*β*_1*−*42_ are from the residues Phe19, Phe20, Val36, Ala21, Val39 and Val40 of A*β*_1*−*42_ and Arg308, Arg296, Lys305, Pro251, Gln330 of HSV-1 gB (see Figure 9). The result of decomposition is similar to the result of visualization in 2.3.

**Figure 9.**
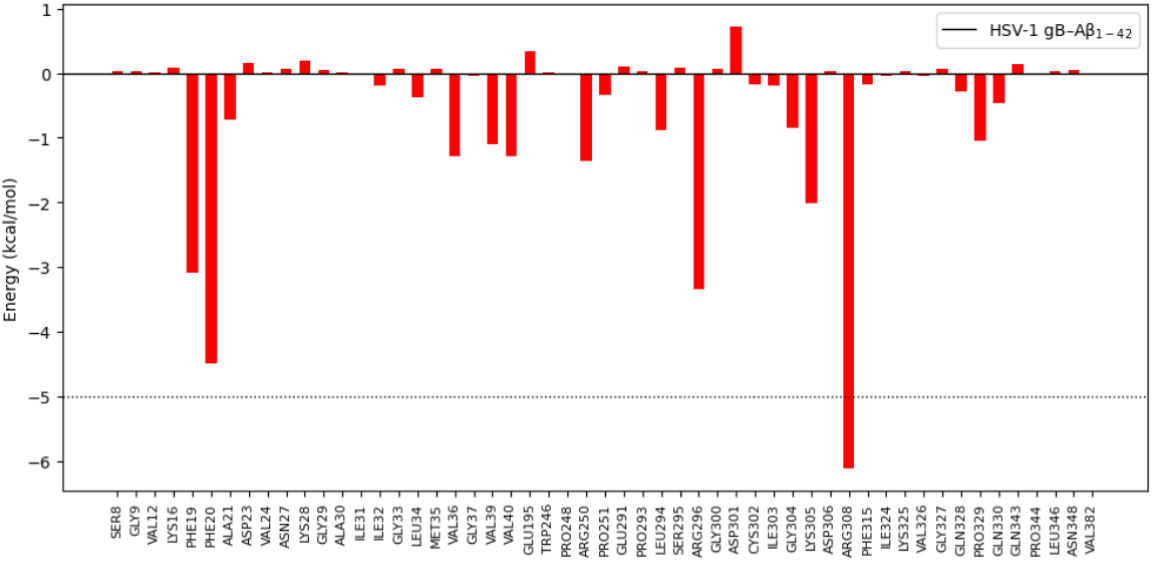
Decomposition of the binding free energy (kcal/mol) on residues of interface for HSV-1 gB – A*β*_1*−*42_ complex.

**Figure 10.**
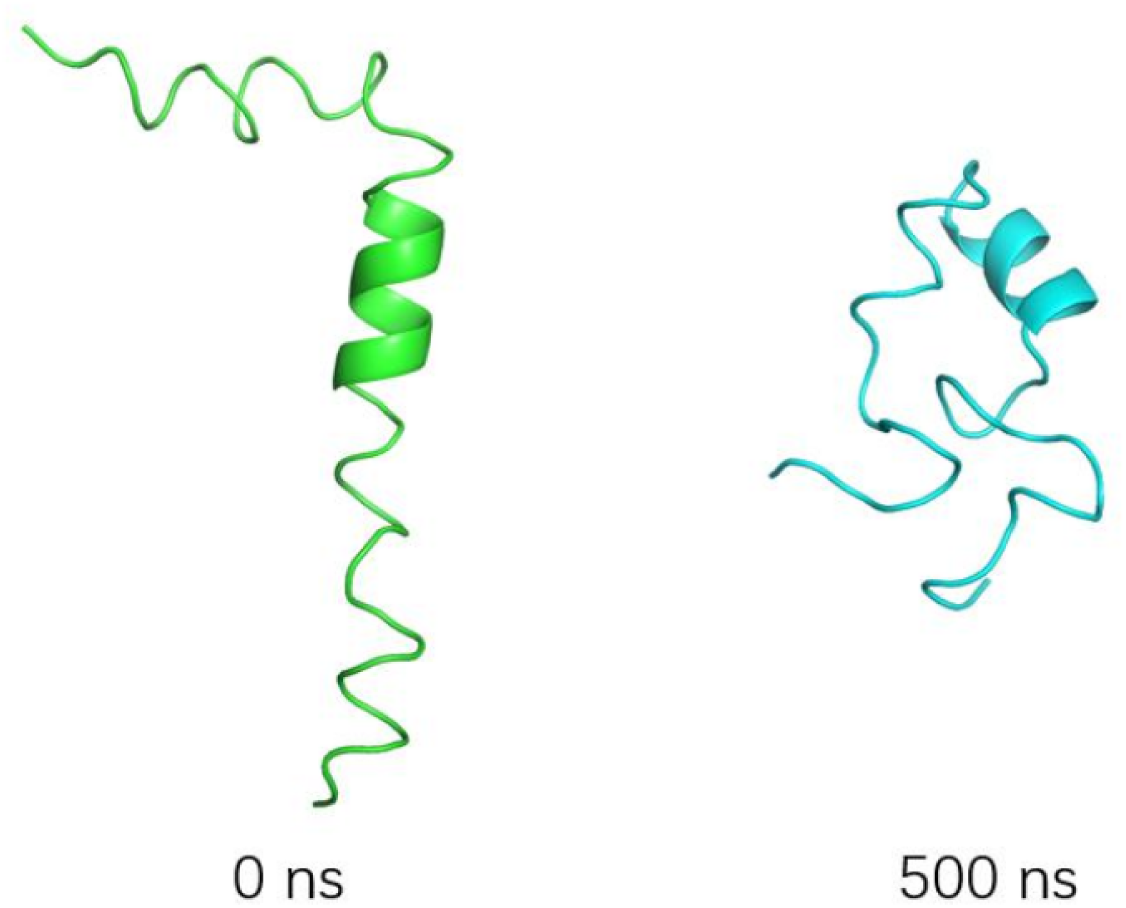
The initial structure (green) and final structure (cyan) of A*β*_1*−*42_ in the 500ns MD simulation.

### A*β*_1_*_−_*_42_ during 500ns MDs

During 500ns MD simulation, a significant reduction in *α*-helix was also observed in A*β*_1*−*42_, similar to what we previously observed in HSV-1 gB –A*β*_1*−*42_ complex, accompanied by a conversion to the *β*-sheet components turn. This conversion ceased after nearly 180ns as shown in Figure 11, and maybe it would continue if we extended the simulation just like Wang [9] had done.

**Figure 11.**
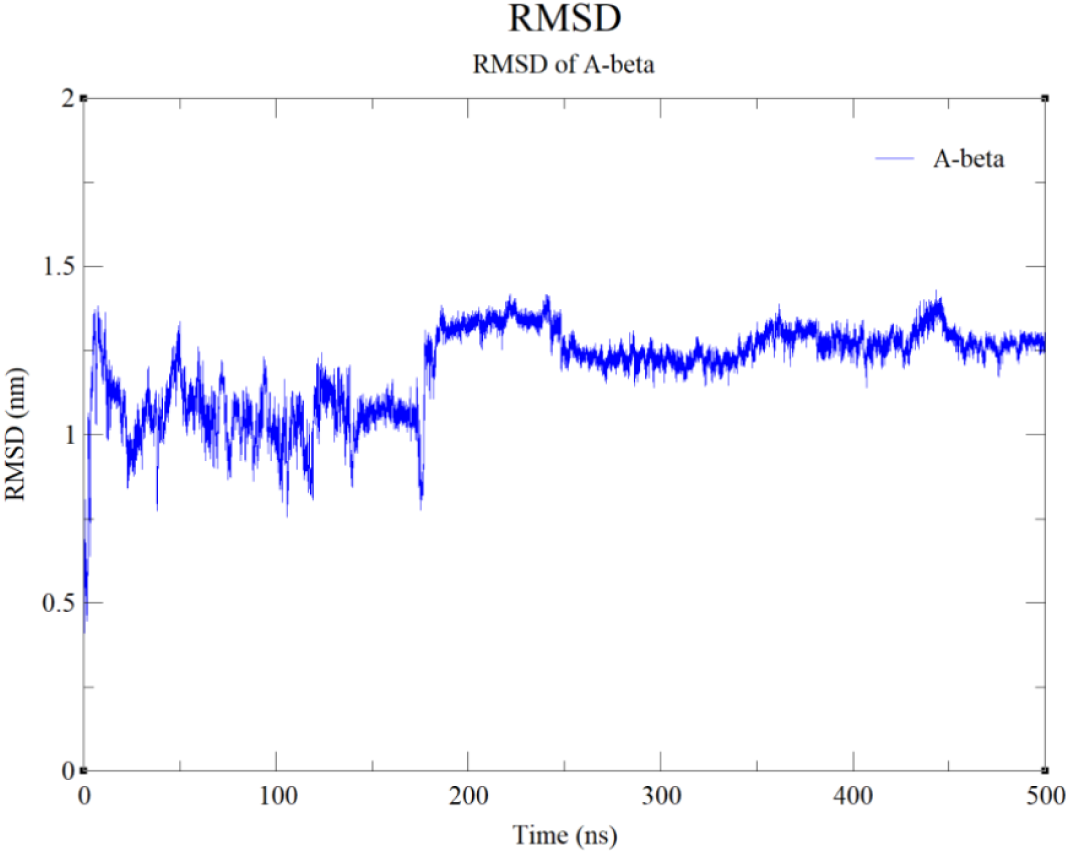
The RMSD of A*β*_1*−*42_ calculated by GROMACS. The conformation of A*β*_1*−*42_ stabilized after nearly 180ns.

### Analysis of secondary structures

Here, we conduct further analysis on the secondary structure of A*β*_1*−*42_ both in the complex and individual simulations during 500ns MD using the timeline plugin contained in VMD. Figure 12 illustrates secondary structural dynamics of all residues in HSV-1 gB – A*β*_1*−*42_ complex. To better observe the conformational conversion in A*β*_1*−*42_, we have provided an expanded view of Figure 12, while Figure 13 focuses specially on the secondary structure changes of A*β*_1*−*42_ in the complex. Consistent with what observed in MD trajectories, a significant reduction in *α*-helix was observed in A*β*_1*−*42_, with a concomitant conversion to the *β*-turn. It is an interesting result that we found the *β*-turn forming, which might interfere the conversion to toxic conformation [1].

**Figure 12.**
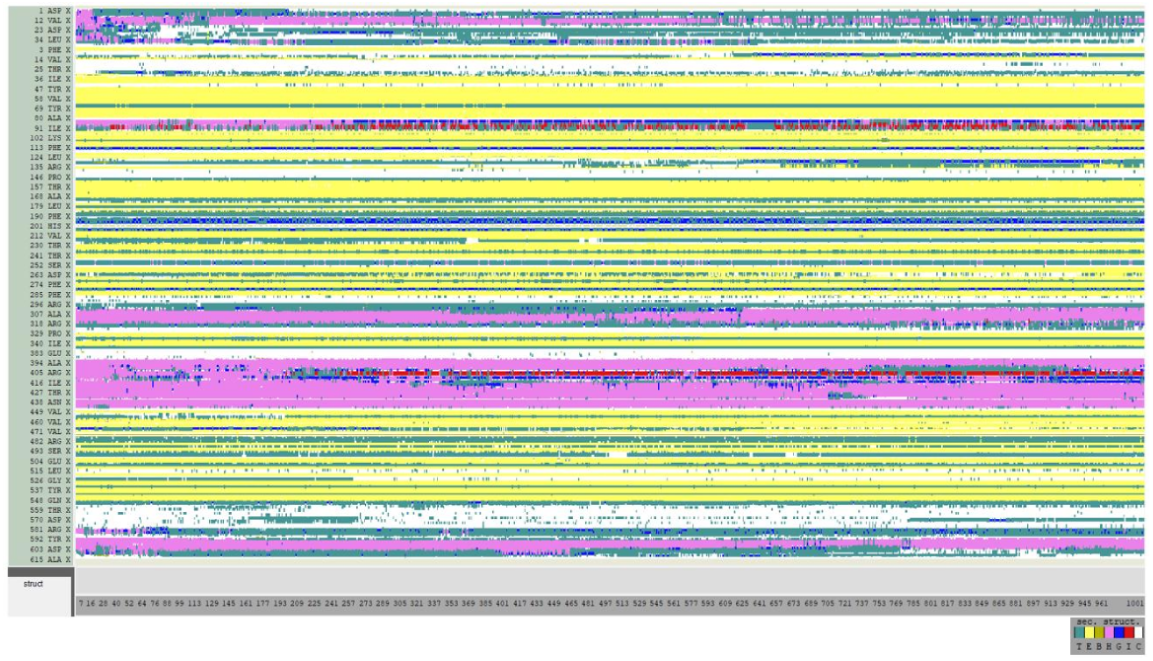
Secondary structure analysis for HSV-1 gB – A*β*_1*−*42_ complex as computed by timeline plugin contained in VMD. The *β*-turn (T) is presented in teal and extended conformation (E) is presented in yellow; isolated bridges are in dark yellow; degrees of helix are in pink (*α*-helix), blue (3-10 helix) and red (*π*-helix); random coils in white.

**Figure 13.**
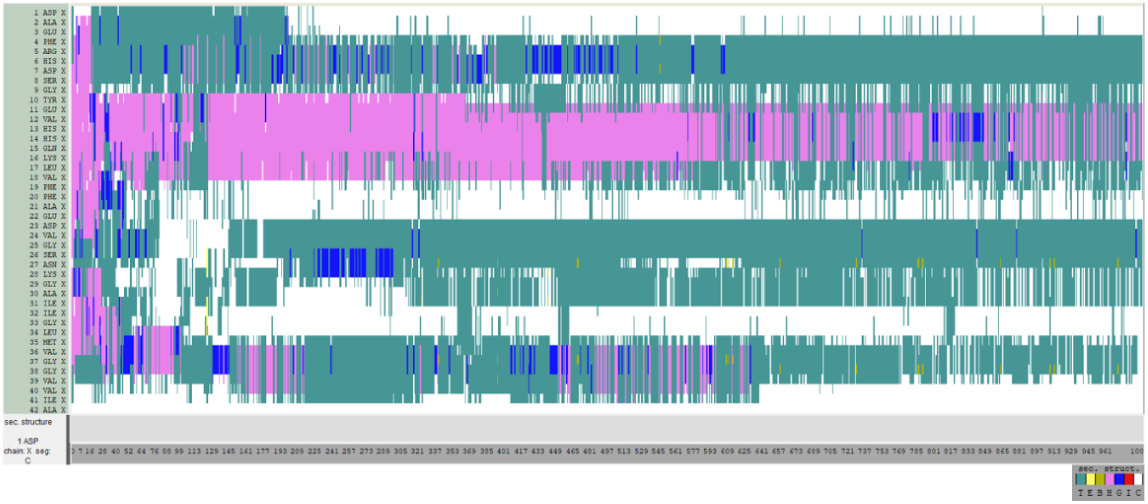
Secondary structure analysis for A*β*_1*−*42_ in HSV-1 gB – A*β*_1*−*42_ complex as computed by timeline plugin contained in VMD. The *β*-turn (T) is presented in teal and extended conformation (E) is presented in yellow; isolated bridges are in dark yellow; degrees of helix are in pink (*α*-helix), blue (3-10 helix) and red (*π*-helix); random coils in white.

The secondary structural dynamics of all residues in A*β*_1*−*42_ during 500ns individual simulation was illustrated in Fig.14. There is also a reduction of *α*-helix, but it occurred from residue No.9 to residue No.16, which is opposite to the one in complex compared with Figure 14.

**Figure 14.**
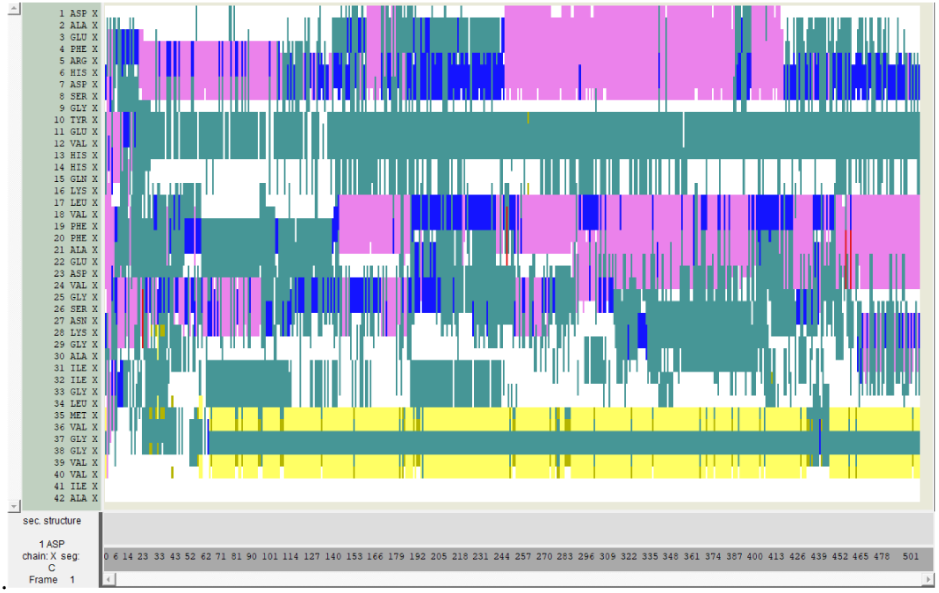
Secondary structure analysis for A*β*_1*−*42_ in the individual simulation as computed by timeline plugin contained in VMD. The *β*-turn (T) is presented in teal and extended conformation (E) is presented in yellow; isolated bridges are in dark yellow; degrees of helix are in pink (*α*-helix), blue (3-10 helix) and red (*π*-helix); random coils in white.

### HSV-gB during 500 ns MDs

The binding site of HSV-gB -A*beta*_1*−*42_ complex was stable during the 500ns simulation. The changes of HSV-gB conformation mostly occurred in its DIII domain (mentioned in Figure 3), which seems irrelevant to our study.

The conventional molecular dynamics analyses presented above demonstrate that the HSV-1 gB–A*β* complex remains structurally stable throughout the simulations while exhibiting changes in residue flexibility, intermolecular interactions, and secondary-structure evolution. We next investigate whether these structural changes are accompanied by modifications in the geometry of the residue interaction network by applying the proposed Forman–Ricci curvature framework.

### HSV-1 Induces localized geometric remodelling

The first objective is to determine whether HSV-1 binding induces persistent changes in the geometry of the residue interaction network. For each residue, the Forman–Ricci curvature was averaged over all frames of the molecular dynamics trajectory for both the isolated A*β* monomer and the HSV-1 gB–A*β* complex.

The resulting average curvature profiles are shown in Figure 15, while the residue-wise curvature differences (complex minus monomer) are presented in Figure 16.

**Figure 15.**
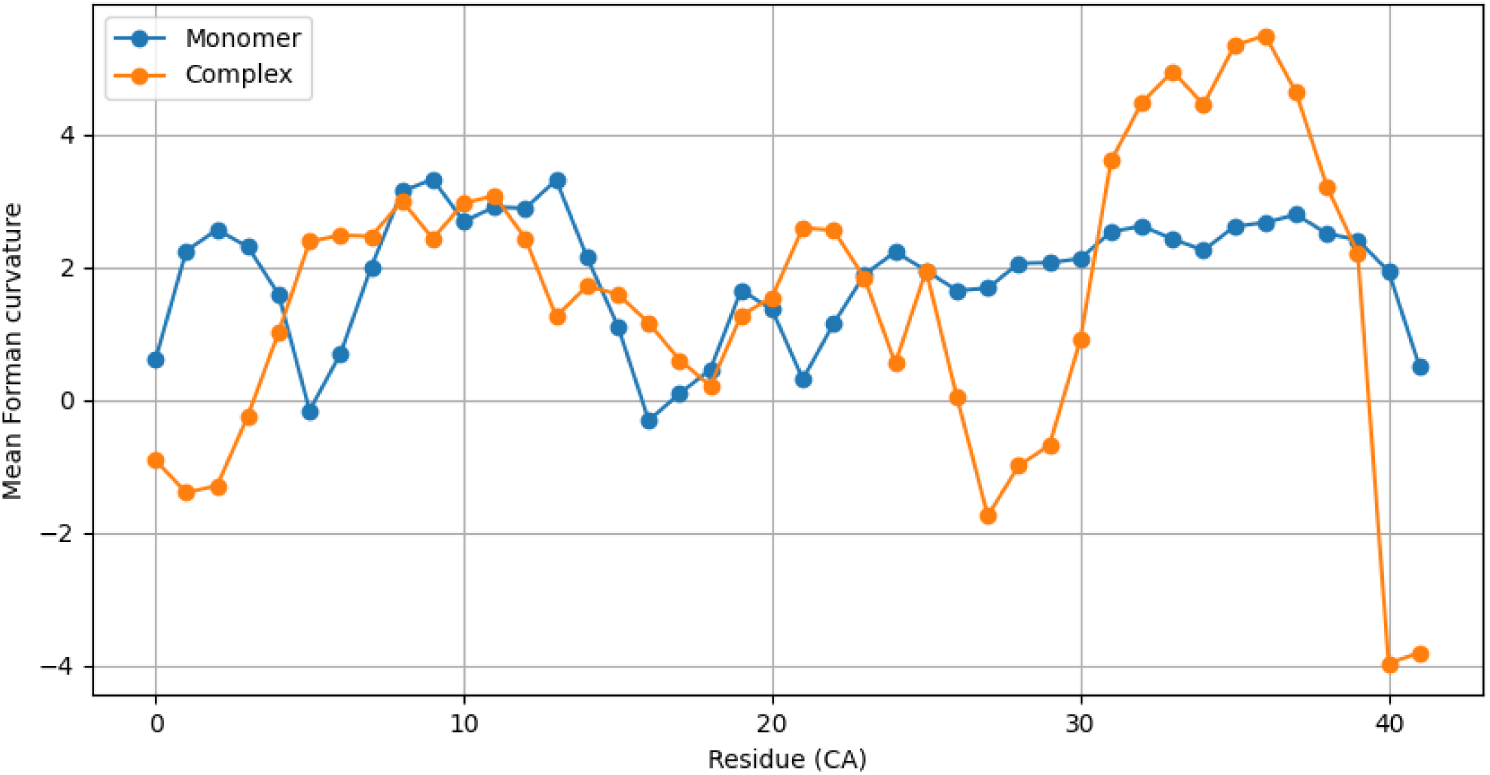
Average Forman–Ricci curvature of every residue throughout the molecular dynamics simulations for the isolated A*β* monomer and the HSV-1 gB–A*β* complex.

**Figure 16.**
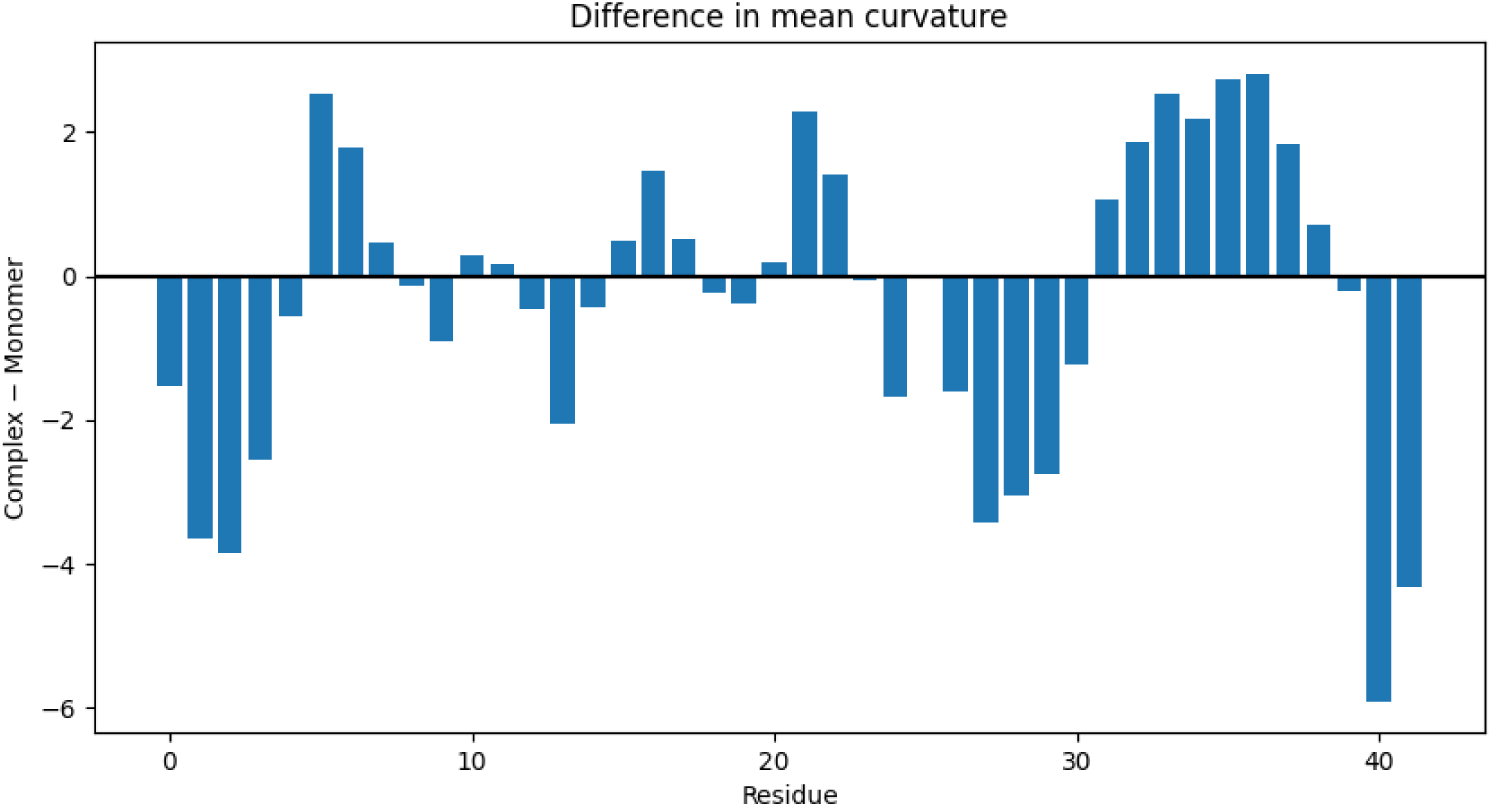
Difference between the average residue curvature of the HSV-1 gB–A*β* complex and the isolated A*β* monomer. Positive values indicate an increase in curvature following viral binding, whereas negative values indicate a decrease.

Although the global average curvature decreases only moderately from 1.89 for the isolated peptide to 1.56 for the HSV-1 complex, this global measure conceals substantial residue-specific geometric remodelling. The distribution of curvature differences spans a wide interval from *−*5.90 to +2.82, while exhibiting a standard deviation of 2.11, approximately six times larger than the mean difference (*−*0.32). This large dispersion indicates that viral binding does not induce a uniform geometric response but instead produces highly localized structural reorganizations.

The largest positive curvature variations are concentrated around residues 33–37, together with residues 5, 6, 16 and 21. In contrast, the strongest negative variations occur primarily within residues 1–3, 27–29 and the C-terminal residues 40–41. Rather than being randomly distributed along the sequence, these changes appear in contiguous residue segments, suggesting that groups of neighbouring residues respond cooperatively to HSV-1 binding.

Interestingly, many of the most strongly affected residues belong to the C-terminal region of A*β*, which is widely recognized as the principal aggregation-prone domain responsible for fibril formation. The localization of the largest curvature variations within this region suggests that the viral protein primarily remodels the geometric organization of residue interactions where aggregation processes are expected to initiate.

Overall, these observations indicate that HSV-1 binding induces pronounced localized geometric remodelling while preserving the overall global geometry of the peptide. Consequently, average Forman–Ricci curvature reveals structural changes that remain largely hidden when considering only global molecular descriptors.

### HSV-1 Stabilises the residue interaction network

While the previous analysis identifies residues undergoing persistent geometric remodelling, it does not indicate whether these new geometric states remain stable throughout the molecular dynamics simulation. To address this question, we quantify the temporal variability of Forman–Ricci curvature for each residue by computing its standard deviation over the entire trajectory. Lower standard deviations correspond to residues whose local interaction networks fluctuate less over time and therefore exhibit greater geometric stability.

Figure 17 compares the residue-wise standard deviations obtained for the isolated A*β* monomer and the HSV-1 gB–A*β* complex.

**Figure 17.**
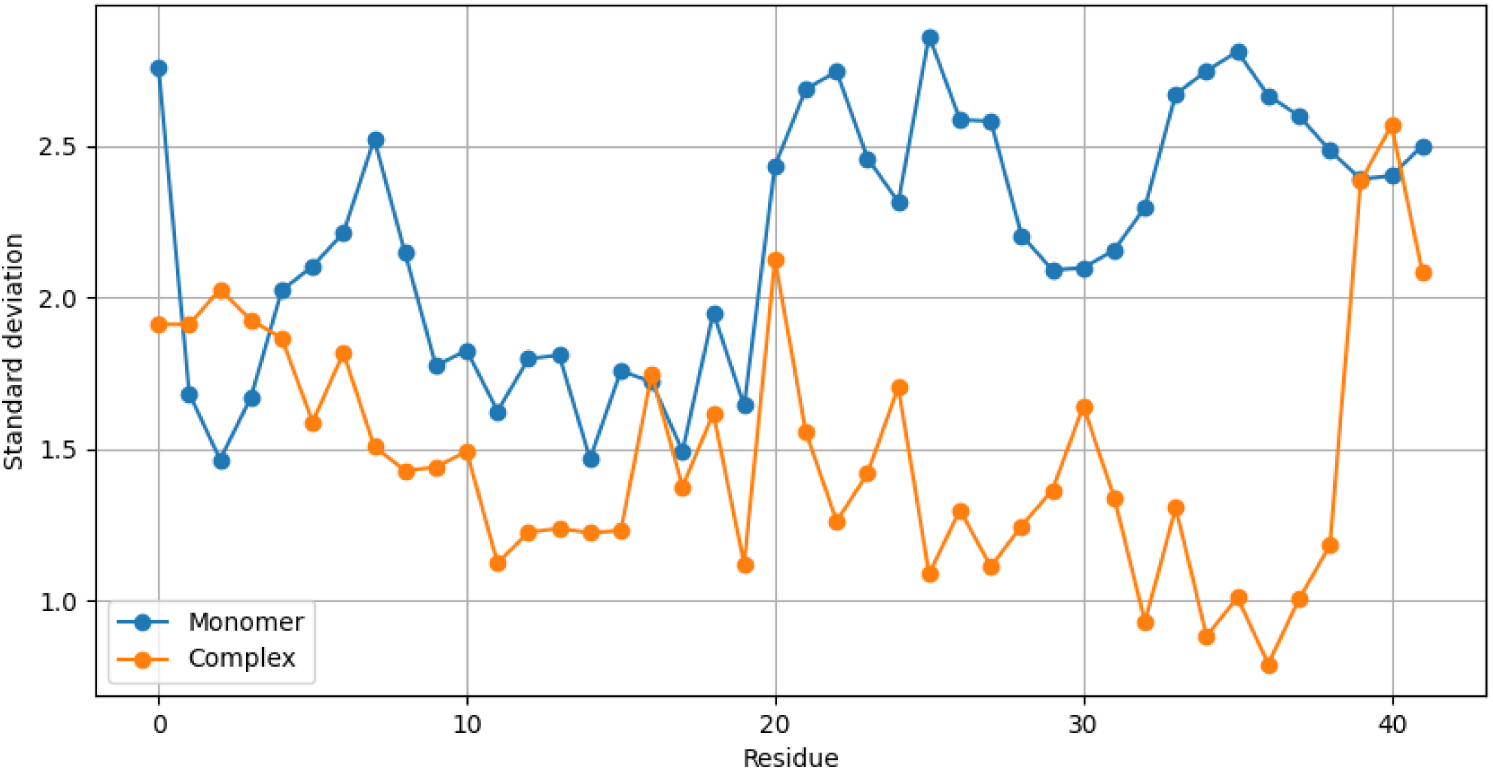
Standard deviation of the Forman–Ricci curvature for each residue during the molecular dynamics simulations of the isolated A*β* monomer and the HSV-1 gB–A*β* complex.

**Figure 18.**
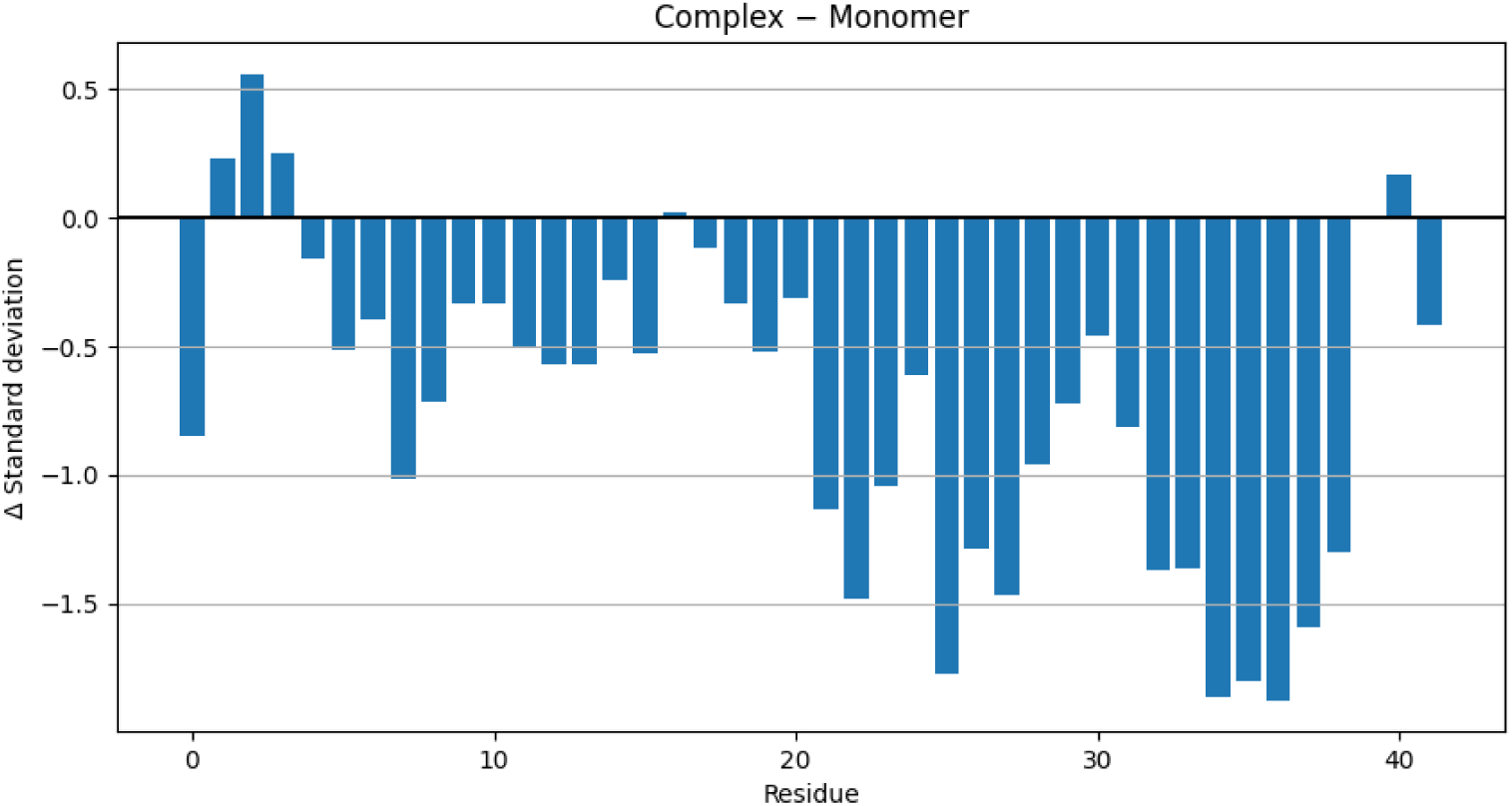
Difference in curvature standard deviation between the HSV-1 gB–A*β* complex and the isolated A*β* monomer. Negative values indicate residues whose geometric fluctuations decrease after HSV-1 binding.

A substantial reduction in curvature variability is observed throughout the peptide following HSV-1 binding. The average residue standard deviation decreases from 2.20 in the isolated monomer to 1.48 in the complex, corresponding to an average reduction of approximately 33%. Furthermore, the mean difference in residue fluctuations is *−*0.72, with only four residues exhibiting increased variability after complex formation.

The largest reductions are concentrated within residues 22–38, particularly around residues 32–37, where the curvature standard deviation decreases by as much as 1.87 units. These residues largely coincide with those identified in the previous section as undergoing the strongest geometric remodelling. This agreement indicates that the regions experiencing the largest structural reorganization are also those that become the most geometrically stable once the complex has formed.

Only a small number of residues located primarily near the N-terminus (residues 1–3) display a slight increase in temporal variability. The maximum increase remains limited to approximately 0.56, which is considerably smaller than the largest reductions observed elsewhere in the peptide. Consequently, the dominant effect of HSV-1 binding is not to increase structural disorder, but rather to stabilize most of the residue interaction network.

Regional analysis further supports this observation. The average fluctuation decreases within all three structural regions of A*β*. The N-terminal segment (residues 0–16) exhibits a reduction from 1.06 to 0.82, the central region (residues 17–29) from 1.26 to 0.68, while the aggregation-prone C-terminal region (residues 30–41) undergoes the most pronounced stabilization, with the average standard deviation decreasing from 1.78 to only 0.68. This nearly threefold reduction indicates that HSV-1 binding strongly suppresses geometric fluctuations within the region most closely associated with amyloid aggregation.

Taken together, these results demonstrate that HSV-1 binding not only remodels the geometry of the residue interaction network but also substantially reduces its temporal variability. The simultaneous observation of localized geometric reorganization (Section 5.1) and decreased fluctuations suggests that viral binding drives the peptide toward more persistent and well-defined geometric configurations, particularly within the C-terminal aggregation domain.

### Geometric remodelling emerges and persists throughout the molecular dynamics trajectory

The analyses presented thus far characterize average geometric properties over the entire simulation. While these global statistics demonstrate that HSV-1 binding induces localized remodelling and increased structural stability, they do not reveal how these geometric differences develop throughout the molecular dynamics trajectory. We therefore investigate the temporal evolution of the Forman–Ricci curvature in order to determine whether the observed remodelling appears immediately after complex formation or progressively emerges during the simulation.

Based on the previous analyses, residues 33–37 were selected for detailed investigation since they consistently exhibited the largest positive curvature variations together with one of the strongest reductions in temporal fluctuations. For each simulation frame, the average Forman–Ricci curvature of these residues was computed and subsequently smoothed using a moving average to emphasize the long-term geometric evolution.

Figure 19 presents the temporal evolution of the average curvature for both molecular systems, while Figure 20 shows the corresponding curvature difference between the HSV-1 gB–A*β* complex and the isolated monomer.

**Figure 19.**
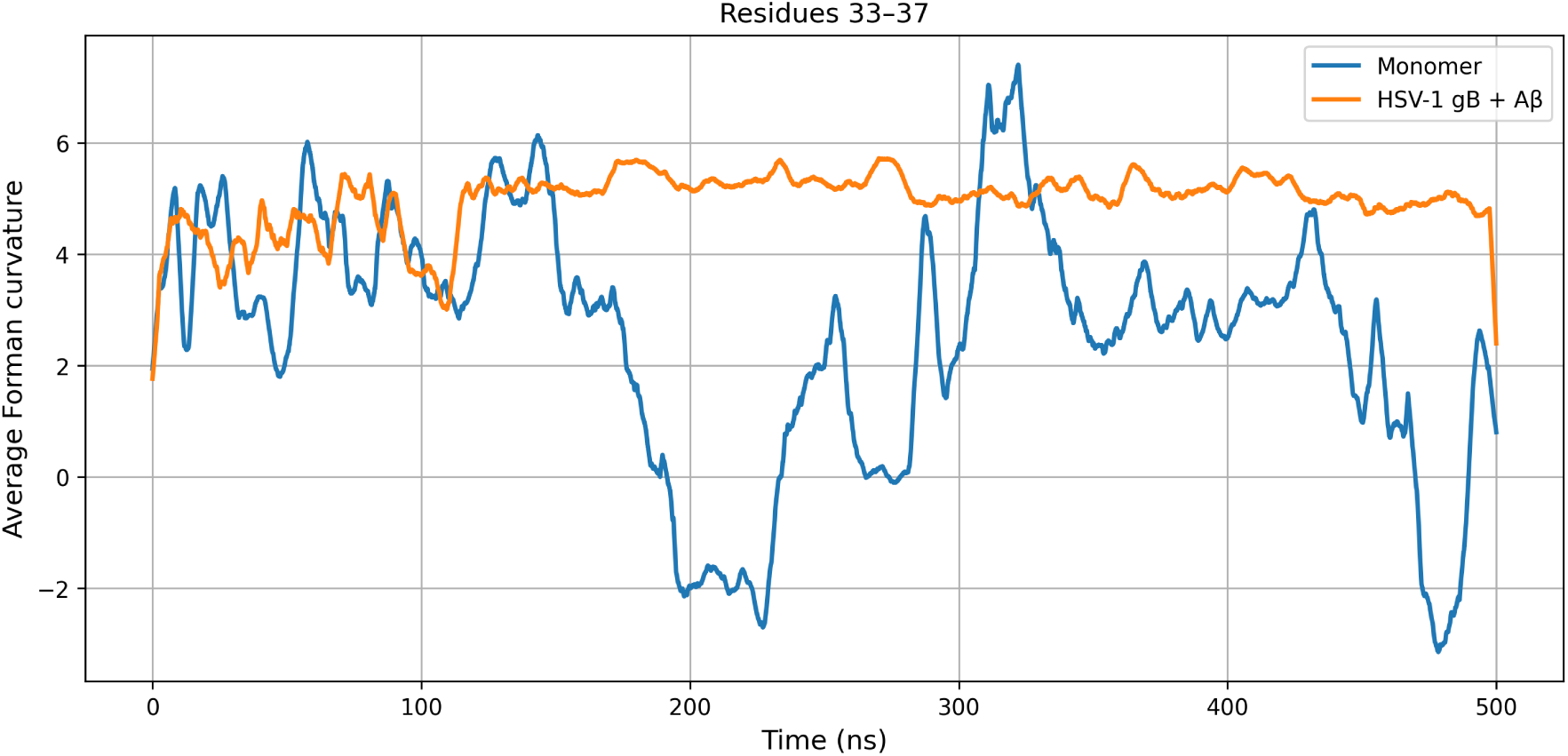
Smoothed average Forman–Ricci curvature of residues 33–37 throughout the molecular dynamics simulations for the isolated A*β* monomer and the HSV-1 gB–A*β* complex.

**Figure 20.**
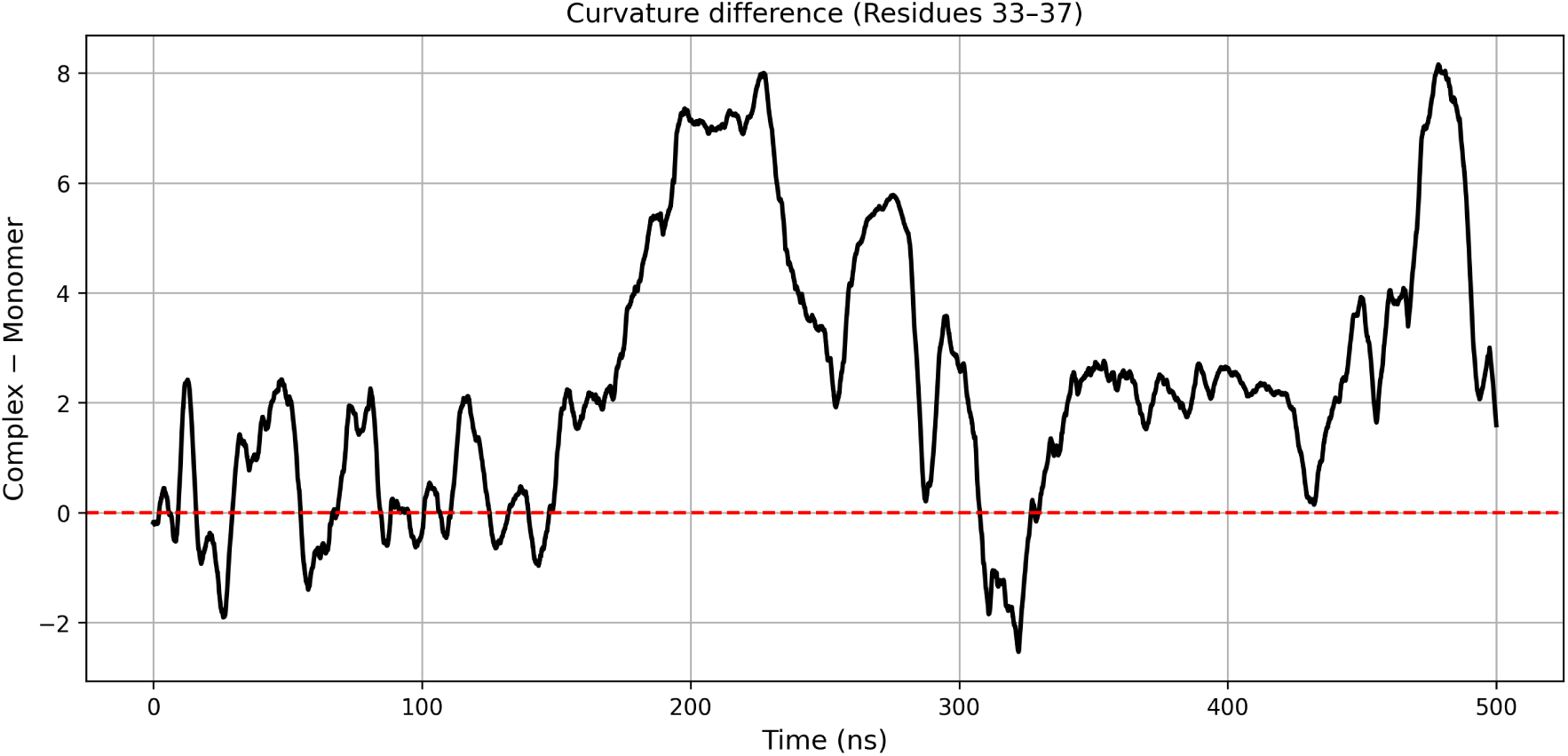
Temporal evolution of the curvature difference between the HSV-1 gB–A*β* complex and the isolated A*β* monomer for residues 33–37. Positive values indicate higher curvature in the complex.

The average curvature of residues 33–37 remains consistently higher in the HSV-1 gB–A*β* complex throughout the entire simulation. The mean curvature difference reaches 2.42, with individual values ranging from *−*5.63 to 11.06. Although short-lived fluctuations are observed, the overall difference remains positive during most of the trajectory, indicating that the geometric remodelling identified in the previous sections is not a transient event but a persistent feature of the bound state.

Interestingly, the average curvature difference increases from 2.19 during the first half of the simulation to 2.65 during the second half. Simultaneously, the temporal variability decreases from 2.93 to 2.37, indicating that the curvature signal becomes progressively more stable as the simulation proceeds. These observations suggest that the residue interaction network gradually converges toward a persistent geometric organization following HSV-1 binding rather than reaching its final state immediately after the simulation begins.

The continuous separation between the monomer and complex trajectories further demonstrates that the observed geometric remodelling is maintained over hundreds of nanoseconds. Rather than representing isolated structural fluctuations, the curvature profiles indicate the establishment of two distinct dynamical regimes: one corresponding to the intrinsic conformational dynamics of the isolated A*β* peptide and the other describing the geometrically remodelled state induced by HSV-1 binding.

These temporal observations reinforce the conclusions of the previous analyses. The C-terminal region not only undergoes the largest geometric reorganization and exhibits the strongest reduction in fluctuations, but also progressively stabilizes into a distinct geometric state that persists throughout the molecular dynamics trajectory. This behaviour strongly suggests that HSV-1 binding modifies the long-term organization of residue interactions rather than merely perturbing local conformations.

### Cooperative geometric domains reveal coordinated structural organization

The previous analyses demonstrated that HSV-1 binding induces localized geometric remodelling, reduces temporal fluctuations, and progressively stabilizes the residue interaction network. However, these analyses treat each residue independently and therefore do not address whether neighbouring residues evolve as isolated entities or as coordinated structural domains.

To investigate the global organization of the geometric response, the complete curvature matrix was analysed throughout the molecular dynamics simulations. Spatio-temporal heatmaps were first constructed to visualize the evolution of Forman–Ricci curvature in both molecular systems. The frame-by-frame difference between the HSV-1 gB–A*β* complex and the isolated monomer was then computed in order to identify regions undergoing persistent geometric remodelling. Finally, hierarchical clustering was performed on the residue curvature trajectories to identify groups of residues exhibiting similar temporal behaviour.

Figure 21 presents the spatio-temporal curvature map of the isolated A*β* monomer, whereas Figure 22 shows the corresponding map for the HSV-1 gB–A*β* complex. The residue-wise curvature differences are illustrated in Figure 23.

**Figure 21.**
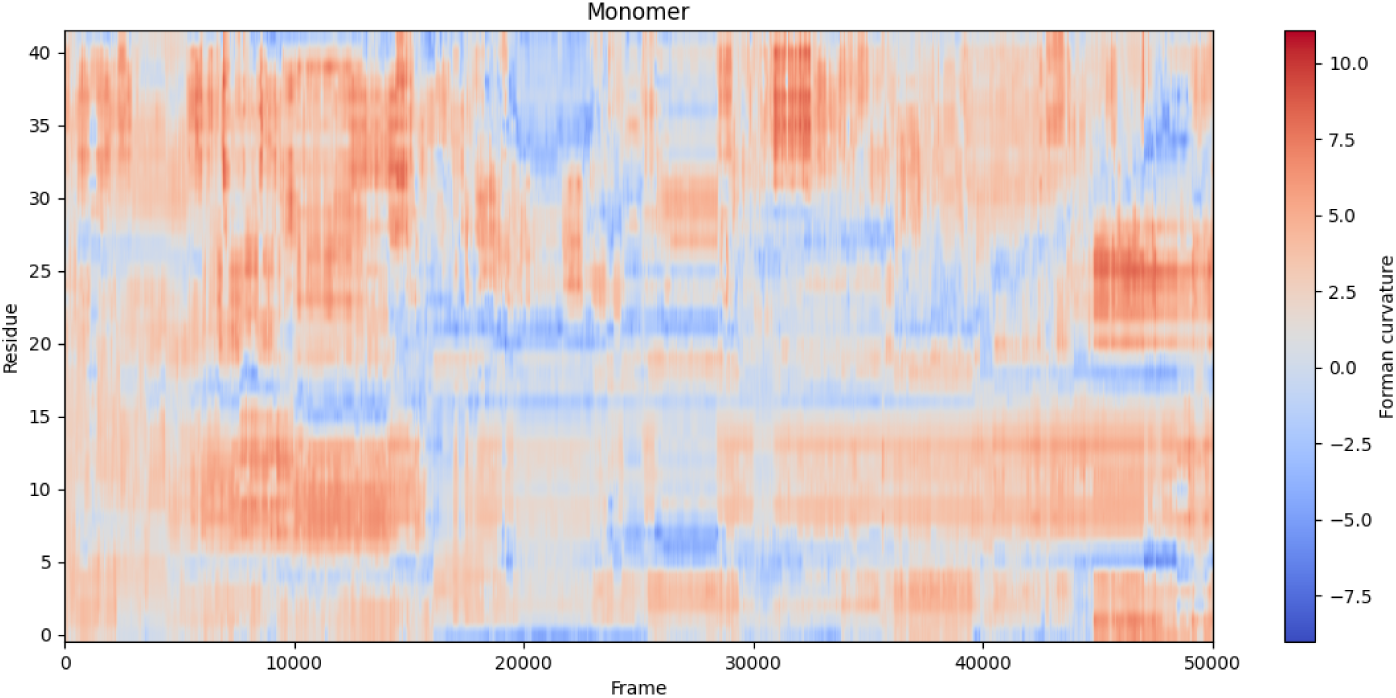
Spatio-temporal distribution of Forman–Ricci curvature for the isolated A*β* monomer.

**Figure 22.**
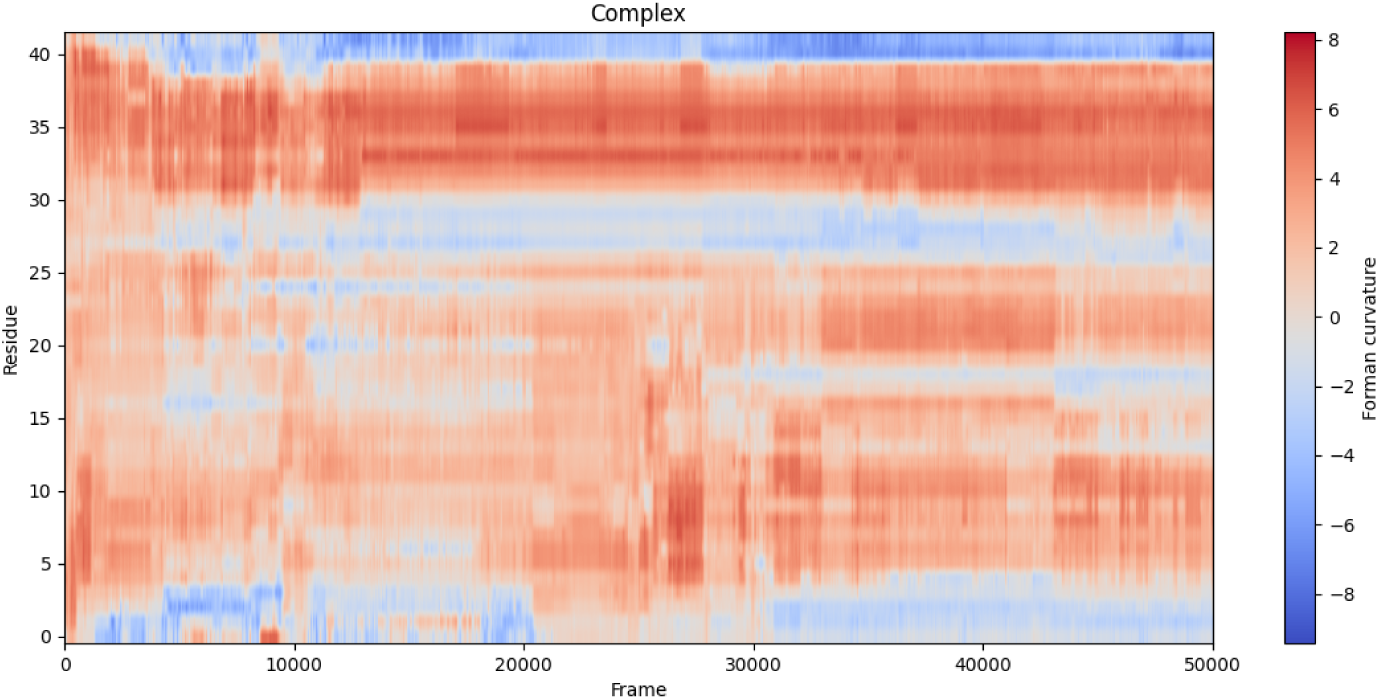
Spatio-temporal distribution of Forman–Ricci curvature for the HSV-1 gB–A*β* complex.

**Figure 23.**
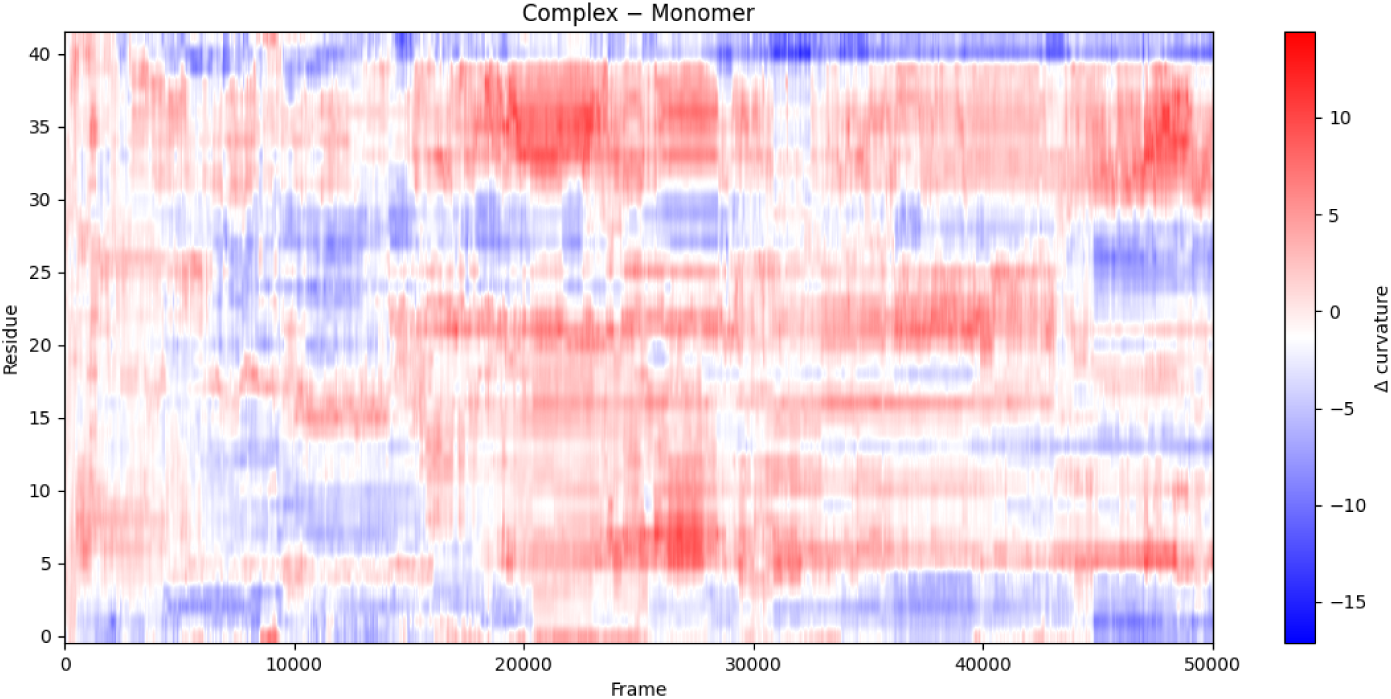
Frame-by-frame difference in Forman–Ricci curvature between the HSV-1 gB–A*β* complex and the isolated A*β* monomer. Positive values indicate higher curvature in the complex.

The heatmaps provide a global view of the geometric evolution of the peptide. Rather than being uniformly distributed along the sequence, the curvature differences are confined to well-defined residue blocks that persist throughout the simulation. This observation confirms that HSV-1 binding induces localized geometric remodelling rather than a homogeneous perturbation of the entire residue interaction network.

Quantitative analysis supports these observations. The largest mean absolute curvature differences are observed for residues 40, 41, 27, 33, 35 and 36, with average deviations exceeding three curvature units. Likewise, residues 40, 39, 0, 7, 6 and 21 exhibit the largest temporal variability in the curvature difference, indicating that these regions experience the most pronounced geometric reorganization during the molecular dynamics trajectory. Importantly, many of these residues were independently identified in the previous analyses, demonstrating the robustness of the observed remodelling.

To determine whether these residues evolve independently or as coordinated structural units, hierarchical clustering was applied to the curvature trajectories of all residues. The resulting dendrogram is shown in Figure 24.

**Figure 24.**
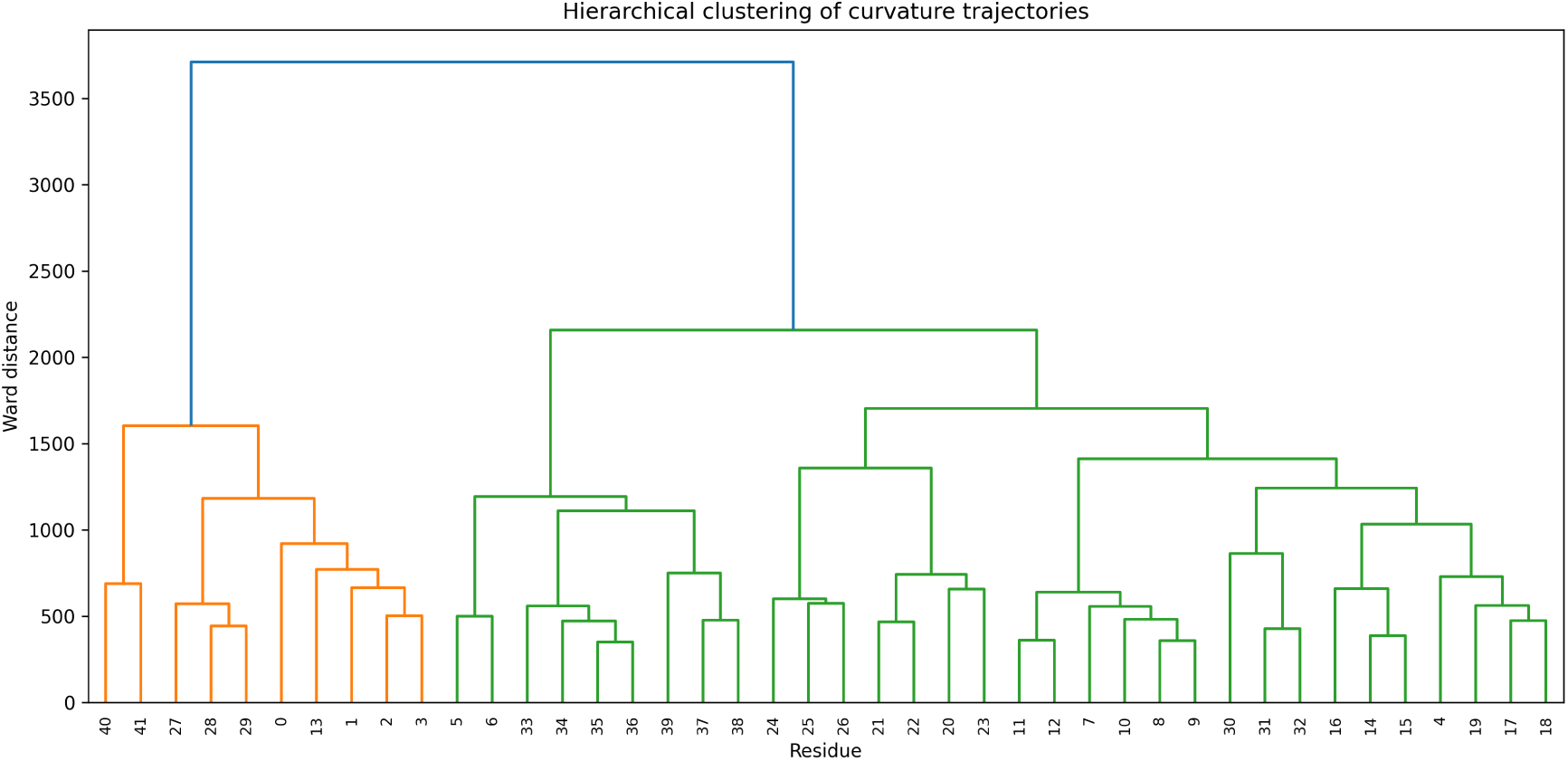
Hierarchical clustering of residue Forman–Ricci curvature trajectories. Residues are grouped according to the similarity of their temporal geometric evolution throughout the molecular dynamics simulations. Four major cooperative geometric domains are identified.

Cutting the dendrogram at the selected linkage threshold reveals four principal geometric domains. Cluster 1 groups residues *{*0, 1, 2, 3, 27, 28, 29, 40, 41*}*, which predominantly exhibit persistent negative curvature variations following HSV-1 binding. Cluster 2 comprises residues *{*32, 33, 34, 35, 36, 37*}* and corresponds to the region displaying both the largest positive curvature increase and the strongest reduction in temporal fluctuations. Cluster 3 includes residues *{*5, 6, 7, 8, 9, 10, 11, 12, 31, 38, 39*}*, which exhibit an intermediate geometric response, while Cluster 4 contains the remaining residues whose curvature profiles remain comparatively moderate throughout the simulations.

Remarkably, the residues exhibiting similar curvature trajectories form contiguous sequence segments rather than isolated amino acids. In particular, Cluster 2 coincides almost exactly with the C-terminal region previously identified as undergoing the strongest geometric remodelling. The emergence of this coherent cluster strongly suggests that HSV-1 binding reorganizes the residue interaction network at the level of entire structural domains rather than through independent local perturbations.

Taken together, the heatmaps and hierarchical clustering demonstrate that the geometric response to HSV-1 binding is highly cooperative. The residue interaction network reorganizes into distinct geometric domains whose temporal evolution remains coherent throughout the molecular dynamics simulation. These results extend the residue-by-residue analyses presented in the previous sections and provide a global description of how viral binding reshapes the geometry of the A*β* interaction network.

## Discussion

The present study combines conventional molecular dynamics analyses with a graph-theoretical description of protein conformational dynamics to investigate the interaction between HSV-1 glycoprotein B and the A*β* peptide. Conventional molecular dynamics analyses establish that the HSV-1 gB–A*β* complex remains structurally stable throughout the simulations, while revealing persistent intermolecular interactions, favourable binding energetics, and conformational changes within the peptide. Building upon these structural observations, the Forman–Ricci curvature analysis provides a complementary description of the same trajectories by characterizing the geometry of the residue interaction network.

Across the four curvature-based analyses, a consistent pattern emerges. HSV-1 binding induces localized geometric remodelling concentrated primarily within the aggregation-prone C-terminal region of A*β*. These regions simultaneously exhibit reduced temporal fluctuations, progressively stabilize throughout the simulation, and organize into coherent cooperative domains identified through hierarchical clustering. The agreement among these independent analyses suggests that the observed curvature patterns reflect robust structural reorganization rather than isolated statistical fluctuations.

Importantly, these geometric observations complement the conventional molecular dynamics descriptors. While RMSD, RMSF, hydrogen-bond occupancy, secondary-structure evolution, and binding free-energy calculations quantify structural stability and intermolecular interactions, Forman–Ricci curvature characterizes how these structural changes reorganize the geometry of the residue interaction network. The two approaches therefore provide distinct but complementary descriptions of the same dynamical process, linking atomic-scale conformational changes to the evolving organization of residue contacts.

The conventional molecular dynamics analyses further indicate that the interaction with HSV-1 gB is accompanied by conformational changes in the A*β* peptide. While isolated A*β* has been reported to undergo a transition from *α*-helical conformations toward aggregation-prone *β*-sheet structures, the HSV-1 gB–A*β* complex predominantly exhibits a transition from *α*-helices to *β*-turns during the timescale explored in the present simulations. Because *β*-turn-rich conformations have been proposed as intermediate states preceding complete *β*-sheet formation, this observation raises the possibility that viral binding modifies the pathway of A*β* conformational conversion rather than simply altering its final structure. Extending the molecular dynamics simulations to longer timescales will be necessary to determine whether these *β*-turn conformations remain stable or ultimately evolve toward fibrillar *β*-sheet assemblies.

Interestingly, the regions undergoing these conformational rearrangements coincide with those identified by the Forman–Ricci curvature analysis as exhibiting the strongest geometric remodelling and the greatest reduction in temporal fluctuations. This agreement suggests that the observed changes in network geometry are not merely mathematical features of the residue interaction graph, but instead reflect biologically relevant structural reorganization associated with HSV-1 binding. Graph curvature therefore provides an additional layer of information linking local conformational changes to the evolving organization of residue interactions.

Beyond the present biological application, this work highlights the broader potential of discrete differential geometry for molecular dynamics analysis. Because graph curvature is computed directly from residue interaction networks, it can be integrated into existing molecular dynamics workflows with minimal computational overhead while providing residue-wise geometric descriptors that naturally support temporal, statistical, and clustering analyses.

Despite these encouraging results, several limitations should be acknowledged. The present framework considers unweighted residue interaction graphs constructed from C*α* contacts and focuses exclusively on Forman–Ricci curvature. Future studies could investigate weighted graph representations incorporating physicochemical properties, alternative notions of discrete Ricci curvature such as Ollivier–Ricci curvature, and larger sets of molecular systems to further evaluate the robustness and generality of the proposed methodology.

## Conclusion

This work presents a graph-theoretical framework for analysing molecular dynamics trajectories through Forman–Ricci curvature and applies it to the interaction between HSV-1 glycoprotein B and the amyloid-*β* peptide. By representing each simulation frame as a residue interaction graph, the proposed methodology complements conventional molecular dynamics analyses with a geometric description of the evolving residue interaction network.

Together, the conventional molecular dynamics analyses and the graph-based framework provide a comprehensive view of the interaction between HSV-1 and A*β*. While the molecular dynamics analyses demonstrate that complex formation is associated with stable binding, favourable energetics, and persistent conformational changes, the Forman–Ricci curvature analysis reveals that these structural changes are accompanied by a pronounced geometric reorganization of the residue interaction network. In particular, HSV-1 binding induces localized curvature remodelling, reduces geometric fluctuations, progressively stabilizes the aggregation-prone C-terminal region, and organizes residues into cooperative geometric domains that remain coherent throughout the simulation.

These findings demonstrate that Forman–Ricci curvature provides information that is complementary to classical molecular dynamics descriptors. Rather than replacing conventional structural analyses, graph curvature enriches their interpretation by quantifying how conformational changes reshape the geometric organization of residue interactions over time.

More broadly, the proposed framework is independent of the specific biological system considered here and can be readily extended to other molecular dynamics studies involving protein–protein, protein–DNA, protein–RNA, or protein–ligand interactions. Because it relies solely on residue interaction graphs, the methodology remains computationally efficient while offering a novel geometric perspective on biomolecular dynamics.

Beyond demonstrating the usefulness of Forman–Ricci curvature for molecular dynamics analysis, the present study also provides new insight into the interaction between HSV-1 and A*β*. The combined structural and graph-based analyses suggest that viral binding reorganizes the residue interaction network while promoting conformational changes within the aggregation-prone C-terminal region of the peptide. Whether the observed *β*-turn-rich conformations represent metastable intermediates on the pathway toward *β*-sheet formation remains an open question that warrants longer molecular dynamics simulations and experimental validation. Addressing this question may further clarify the molecular mechanisms by which HSV-1 influences A*β* aggregation and Alzheimer’s disease pathology.

An important future direction is the integration of graph curvature with Topological Data Analysis. Whereas Forman–Ricci curvature characterizes the local geometry of residue interaction networks, persistent homology captures their global topological organization across multiple spatial scales. Combining these complementary mathematical frameworks could provide a unified geometry–topology description of biomolecular dynamics, enabling the simultaneous characterization of local geometric remodelling and global topological evolution during molecular recognition and conformational transitions.

## Acknowledgments

This research is supported by the grant PID2023-146683OB-100 funded by MICIU/AEI /10.13039/501100011033 and by ERDF, EU. Additionally, it is supported by Ikerbasque Foundation and the Basque Government through the BERC 2026-2030 program and by the Ministry of Science and Innovation. It is also supported by the French National Research Agency (ANR) under grant ANR-25-CE45-1521.

